# DruID: Personalized **Dru**g Recommendations by **I**ntegrating Multiple Biomedical **D**atabases for Cancer

**DOI:** 10.1101/2021.04.11.439315

**Authors:** Herty Liany, Anand Jeyasekharan, Vaibhav Rajan

**Affiliations:** National University of Singapore

## Abstract

Advances in next-generation sequencing technologies have led to the development of personalized genomic profiles in diagnostic panels that inform oncologists of alterations in clinically relevant genes. While targeted therapies for some alterations may be found, an effective therapeutic strategy should consider multiple and dependent genetic interactions that affect cancer progression, a task which remains challenging. There are ongoing efforts to profile cancer cells in-vitro, both to catalog their genomic information and study their sensitivity to various drugs. There is a need for tools that can interpret the personalized genomic profile of a patient in light of information from these biological and pre-clinical studies and recommend potentially useful drugs. To address this need, we develop a new algorithmic framework called DruID, to effectively combine drug efficacy predictions from a deep neural network model with information, such as drug sensitivity, drug-drug interactions and genetic dependencies, from multiple publicly available databases. We empirically evaluate DruID on cancer cell line data on which efficacy of many drugs have been experimentally determined. We find that DruID outperforms competing approaches and promises to be a useful tool in clinical decision-making.

## 1 Introduction

### 1.1 Objective

Advances in next-generation sequencing technologies have led to the development of personalized genomic profiles that can now be used in clinical diagnostics. These diagnostic panels inform oncologists of alterations in specific genes, for which targeted therapies are available or being investigated. However, effectively utilizing the information in these panels to guide treatment decisions remains difficult due to various reasons [1, 2]. For example, therapies targeting specific genetic alterations may not always be effective, and the cancer may become resistant to them [3, 4, 5]. Suitable combination of therapies that target multiple alterations or that combine targeted therapies with traditional chemotherapy, may need to be found which can also be challenging [6, 7].

Genetic alterations and their dependencies are not fully understood and there are several ongoing efforts to profile cancer cells in-vitro, both to catalog their genomic information and to study their sensitivity to various drugs [8, 9, 10]. There is a need for tools that can integrate information from these biological and pre-clinical studies to analyze and interpret the personalized genomic profile of a patient and recommend potentially useful drugs [11]. To address this need, we aim to develop an algorithmic framework called **DruID** (**Dru**g Recommendations by **I**ntegrating Multiple Biomedical **D**atabases), to effectively combine a deep neural network model for predicting drug efficacy on a genomic profile with information from multiple databases, that can include drug sensitivity information on model cancer cell lines, drug-drug interactions and genetic dependencies such as synthetic lethality.

### 1.2 Background and Significance

Cancer treatment is challenging due to inter-tumor and intra-tumor heterogeneity that results in considerable diversity in subtypes, treatment sensitivity and outcomes across patients. As a result, cancer care has been progressively moving from a ‘one-size-fits-all’ approach to a more personalized strategy based on patient-specific molecular characteristics [12]. Such personalization has been enabled by *targeted therapies* – drugs designed to interfere with a specific molecular target, typically a protein, that is believed to have a critical role in tumour growth or progression [13]. These therapies have mainly focused on targeting recurrent actionable mutations in oncogenic drivers. Yet, many challenges remain due to heterogeneity in genomic alterations across tumors, low prevalence of some alterations and emergence of acquired resistance [6, 5]. Clinical significance of many of the alterations are unknown and the list of actionable cancer driver genes remains incomplete [14, 1, 15]. Suitable drugs to target specific alterations may not be known [16], cancers may harbour loss-of-function mutations in tumour-suppressor genes or activating mutations in currently undruggable oncogenes [17], or the right treatment approach may require a combination of drugs that may be difficult to find [6].

The dynamic changes in the genome due to cancer lead to various genetic aberrations such as somatic mutations, copy number variations, changed gene expression profiles, and different epigenetic alterations. There has been concerted efforts to create many public databases to catalog these molecular phenotypes. For instance, The Cancer Genome Atlas (TCGA) [18] has data from more than 11,000 patient across 33 cancer types. Cancer cell lines derived from naturally occurring tumours have been generated and the effect of many drugs on them have been documented, e.g., in the Genomics of Drug Sensitivity in Cancer (GDSC) database [8] and Cancer Cell Line Encyclopedia [9], to aid therapeutic development. The analysis of such molecular and genomic data has vastly improved our understanding of the underlying mechanisms of cancer [19]. Advances in next-generation sequencing technologies have also led to the development of customizable gene panels that have reached the clinic. For instance, Foundation Medicine provides FDA-approved genomic profiling tests comprising clinically relevant biomarkers and genomic alterations which can potentially be used by oncologists to match the patient to appropriate targeted therapies [20].

While target drugs for each alteration may be found, the efficacy of drugs for a given combination of mutations is difficult to determine. This restricts the applicability of targeted therapies to a ‘one target - one drug’ mode which may have limited efficacy. Cancer progression occurs through accumulation of multiple interacting genetic interactions [21, 22], and so, the entire mutational landscape should be considered for making treatment decisions. Thus, for oncologists to utilize these diagnostic profiles, there is a need for tools that can analyze the collective information of the alterations found to determine the therapeutic strategy. Such tools can leverage the growing catalog of information in public databases to learn latent correlations among alterations, including those that are not oncogenes, and integrate known pharmacological evidence, to recommend drugs personalized for the input profile.

Previous studies have attempted to provide such recommendations through predictive models that can predict the efficacy of a drug for an input genomic profile. For instance, [23] proposed an algorithm based on combined drug and cell line similarity score to prioritize drugs specific to a cell line (or a patient profile) based on similarity of the input drug and cell line to those in GDSC and CCLE. Recently, [24] design a neural network that takes as input mutation and expression data from cell lines to predict drug responses. These methods require gene expression data as inputs which are less commonly used in clinical settings. To predict drug response from mutation profiles alone, [25] develop DrugCell, an interpretable deep neural network that takes as input mutation data (from a cell line) and drug chemical structure to predict drug response. Its interpretability comes from a hierarchical network architecture that is based on the structure of biological processes from the Gene Ontology (GO) database. However it is difficult to extend its use across panels because its network architecture would have to change for different diagnostic panels containing different sets of genes, based on the GO hierarchy for the input genes.

To our knowledge, PanDrugs [26] is the only previous method that integrates evidence from multiple public databases and can be utilized to prioritize drugs according to individual genomic data. Pandrugs collects pharmacological data and drug annotations from 24 databases comprising a variety of clinical, biological, and pharmacological sources. Drug names are standardized and annotated with additional information such as drug indication status and gene-drug relationships. PanDrugs also provides two scores that are calculated by integrating information from these databases. Gene Score (GScore) measures the biological relevance and clinical implication of the gene in cancer. Drug Score (DScore) is an estimate of drug response and treatment suitability with respect to a gene. PanDrugs has a web interface where a list of mutated genes (optionally, with variant information) can be given and a list of drugs, prioritized based on their GScore and DScore, can be viewed.

In this paper we present a new, general framework, called DruID, to utilize both predictive models and public databases for drug recommendations personalized to a input genomic profile. A Prescriptive Analytics framework based on Integer Programming is used within DruID to integrate information from diverse sources. Previous approaches such as drug efficacy predictors and precomputed scores like GScore and DScore can be effectively combined through DruID as we demonstrate in our experiments.

## 2 Materials and Methods

We first provide a brief overview of the publicly available data used in DruID. Then we give a detailed description of how DruID integrates the available information.

### 2.1 Data

We describe the publicly available datasets that have been integrated using DruID and used in our experiments.

- **Cancer Cell Lines**. The Cancer Cell Line Encyclopedia (CCLE) [27, 9] is a collection of more than 1800 human cancer cell lines along with pharmacological profiling of 24 drugs on 504 of these cell lines. Genomic characterization of the cell lines have been performed, using various platforms, to obtain somatic mutation profiles, DNA copy numbers, DNA methylation, RNA expression, protein expression and other data. The Genomics of Drug Sensitivity in Cancer (GDSC) database [8] also contains drug sensitivity data of nearly 450 drugs on 1000 cancer cell lines. The data is linked to genomic data (somatic mutations in cancer genes, gene amplification and deletion, tissue type and transcriptional data) to facilitate biomarker and drug discovery. CCLE and GDSC share 988 cell lines in common. Pharmacological profiles have been obtained by exposing cells to different concentrations of a drug and scoring cell viability. The viability relative to untreated controls is typically assumed to follow a sigmoid response as a function of the logarithm of the drug concentration. Thus, a sigmoid curve is fitted to the experimental dose response data to estimate a non-linear mixed effect model. The Area Under the Curve (AUC) is a measure of the efficacy of the drug: small AUC values indicate strong response, whereas large values indicate limited or no response [28]. Thus, AUC value is a combined single value representing potency and efficacy of drug in inhibiting the growth in cell line. More details can be found in [8, 29, 30].
- **Drug-drug Interaction.** Drug interactions are important to consider in combination therapy regimens to maximize efficacy of their synergistic use and minimize toxicity due to antagonistic interactions [7]. The Therapeutic Target Database (TTD) [31] contains detailed information of drugs and their targets, such as target function, sequence and 3D structure, and drug structure, therapeutic class, and clinical development status. Drug Combination Databases (DrugCombDB [32] and DCDB [33]) contain information of synergistic and antagonistic combinations of drugs, curated from multiple sources (e.g., the Food and Drug Administration (FDA) electronic orange book and clinical studies from the literature).
- **Synthetic Lethality.** A pair of genes is considered Synthetic Lethal (SL) when the cell remains viable with functional loss of either gene but the loss of *both* genes is lethal. SL holds great promise for developing targeted anticancer therapies [17]. The key idea is to exploit the presence of an SL pair (A,B) where one of them (say, A) may be mutated in cancer cells. Then, a drug targeting B would kill cancer cells but normal cells, with functional A, would remain viable even with the loss of B. This would lead to highly specific therapies with minimal side effects. Thus, SL can be leveraged to identify novel drug targets in cancers driven by loss of a tumour-suppressor gene or a currently undruggable oncogene and can expand the scope of precision medicine [17, 34]. SynLethDB [35] provides a database for human SL gene pairs, in turn collected from various sources such as from shRNA and RNAi screens [36], the DECIPHER project [37], BioGRID [38], DAISY [39] and text mining. In total there are 19,952 predicted human SL pairs.
- **PanDrugs Scores.** PanDrugs provides two scores – Drug Score (DScore) and Gene Score (GScore) – that are precomputed and stored in their database. DScore measures the therapeutic suitability of a drug with respect to a particular gene (alteration). The computation of DScore takes into account multiple factors – drug-cancer type indication, drug clinical status, in-vitro drug screening data, gene–drug relationship and number of curated databases supporting that relationship. GScore measures the biological relevance of the gene in the tumoral process and its therapeutic actionability. The computation of GScore considers genomic feature evidence, relevance in cancer estimated by various cancer data portals, gene essentiality and its clinical implications. More details can be found in [26]. Note that the computation of these scores utilizes information from multiple public databases.

The datasets obtained from these sources required minimal preprocessing as described in the following before their use in DruID. For SL pairs from SynLethDB, we choose a threshold value (*t* = 0.9): all pairs with scores above *t* are considered SL and given value 1, others are given value 0. Further, we consider only those pairs that have been validated from genomeRNAi experimental screens to minimize false positives. We add 1 to the Drug Interaction scores obtained from the database to make the range [0, 2]. Thus, 0 indicates antagonistic interaction, 1 indicates no (known) interaction and 2 indicates synergistic interaction: higher values are more favorable. For drug efficacy we use (1-AUC) values: thus higher values indicate better drug efficacy on the cell line. The ordering of entities (genes, drugs and cell lines) is ensured to be uniform across all datasets.

### 2.2 Our Model: DruID

DruID takes a list of genes with mutations, from a single patient or cell line, as input. This list may be obtained from genomic profiles such as the FoundationOne diagnostic test [20]. Using multiple public databases, DruID aims to find a list of drugs most suitable for the input set of genomic alterations. DruID is an Integer Programming model that is designed to simultaneously find (i) a set of cell lines, (ii) a set of genes in the cell lines and (iii) a set of drugs, in a manner such that multiple criteria are satisfied. There are two input parameters *L* and *M* to specify the minimum number of cell lines to select and maximum number of drugs. The criteria used are as follows.

- DruID selects a set of at least *L* cell lines and a set of mutations such that:

– the selected cell lines contain all the input mutations, thereby selecting the “closest” set of cell lines similar to the input sample.
– the GScores of selected mutated genes in the cell lines are maximized. Note that these genes need not be just those in the input set.
- DruID selects a set of at most *M* drugs such that:

– the sensitivity of the selected drugs to the selected cell lines is maximized.
– synergistic interactions among the selected drugs is maximized and antagonistic interactions are minimized.
– efficacy (measured by DScores) on SL partners of input genes is maximized.
– predicted drug efficacy (using a predictive model) on the selected cell lines is maximized.

The inputs to our model are summarized in Table 1, along with the notations used in the following. More details of the deep neural network used as the predictive model are in section 2.2.2.

**Table 1:** Input to DruID: matrices from public databases and predictive model

| Matrix (Symbol) | Row (index) | Column (index) | Datatype Description | Source |
| --- | --- | --- | --- | --- |
| 1. Cell line - Drug efficacy (C) | Cell line (i) | Drug (k) | Float, (1-AUC), range: 0–1 | CCLE, GDSC |
| 2. Cell line – Gene mutations (Q) | Cell line (i) | Gene (j) | Binary, 1: damaging non-conserving mutation | CCLE |
| 3. Drug-Drug Interaction (N) | Drug (k) | Drug (k) | Values: {0,1,2}.<br>0: antagonist, 1: no interaction<br>2: synergistic | TTD, DCDB, DrugCombDB |
| 4. DScore (D) | Gene (j) | Drug (k) | Float, range: 0–1 | PanDrugs |
| 5. GScore (G) | Gene (j) | – | Float, range: 0–1<br>(one score per gene) | PanDrugs |
| 6. SL Gene Pair (S) | Gene (j) | Gene (j) | Binary<br>1 if gene pair is predicted/<br>known to be SL | SynLethDB |
| 7. Predicted Drug Efficacy (R) | Drug (k) | – | Prediction for each drug on input,<br>Float, range: 0–1 | Neural Network |

#### 2.2.1 DruID Integer Programming Model

We define the following binary decision variables whose values are learnt through the optimization:

- *z_k_*: to select drugs, *k* = 1,…, *K*
- *t_i_*: to select cell lines, *i* = 1,…, *I*
- *e_j_*: to select genes, *j* = 1,…, *J*

We now describe the objectives and constraints in the model.

##### Cell Line Drug Efficacy

The first objective is to select those cell lines and drugs such that the efficacy of the selected drugs on the selected cell lines is maximized. This is done through the following objective term.

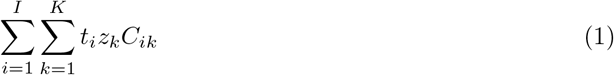

##### GScores

We maximize the GScores of the selected genes in the selected cell lines:

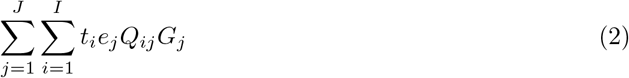

##### Cell Line Selection

We minimize the number of cell lines selected by maximizing:

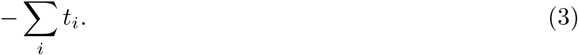

Without this term, the optimization can yield trivial solutions by selecting all the input cell lines.

##### Drug-Drug Interactions

To maximize the synergistic effect (and minimize antagonistic effect) of the selected drugs, we maximize:

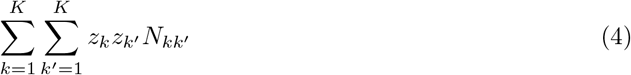

##### SL Interactions

To identify SL partners of selected genes, we introduce another binary indicator *W_j′_* that is set to 1 if gene *j′* is an SL partner gene to one or more of input genes. This can be done by iterating through the SL matrix *S* and checking if there are reported partner genes for any of the input genes. Note that *W* is an input to and not a variable in the optimization model. We maximize the DScores of the selected drugs on the partner genes through:

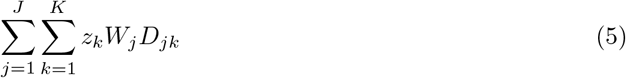

##### Predictive Model

To maximize the predicted drug efficacy on the input cell line/patient, we maximize:

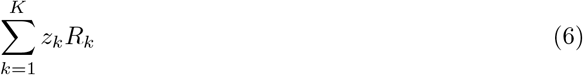

Predictions from any model, e.g., logistic regression, decision trees or neural network, may be used. More details of the neural network we use are in section 2.2.2.

##### Mutation Coverage Constraints

We ensure that at least one cell line is present for each selected mutation, this ensures that the selected cell lines contain all the mutations. This is enabled through the standard IF-ELSE construct: if *e_j_* = 1 then 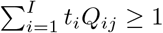. We introduce a large constant *B*_2_ and a binary variable *v_j_*. The following two sets of constraints implement the IF-ELSE construct.

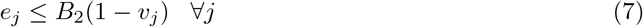

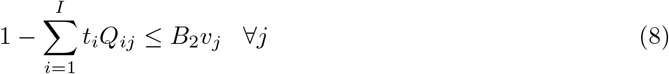

##### Input Mutation Constraints

Further, we ensure that all the input mutations are covered:

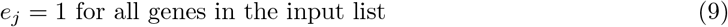

##### User Input Constraints

To ensure at most *M* selected drugs and at least *L* selected cell lines, we add the constraints:

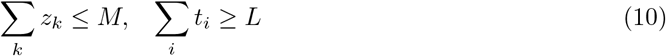

Thus, we get the following Integer Programming Model, with objective terms from (1), (2), (3), (4), (5) and (6). We add the denominators to ensure that each term is normalized and lies between 0 and 1. Weights *α, β* can be set by the user to increase the relative importance of the terms corresponding to the known drug efficacy on selected cell lines and predicted drug efficacy on input cell line.

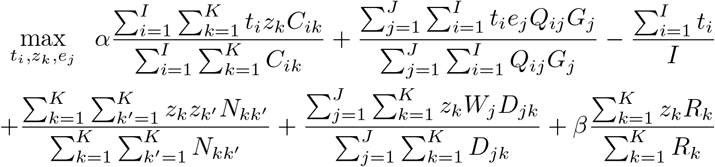

subject to Mutation coverage constraints (7) and (8), input mutation constraint (9) and user-selected maximum drugs and minimum cell lines constraints (10). Figure 1 shows a schematic of DruID.

Integer Programming models, including DruID, can leverage accurate and efficient heuristics, such as LP-based Branch and Bound and Cutting Plane methods, with readily available implementations in general-purpose solvers (e.g. Gurobi [40]) to find approximate solutions. These heuristics are designed to run for a pre-specified period of time, during which they can find multiple optimal and near-optimal solutions.

#### 2.2.2 Deep Neural Network

A predictive model is used in term (6) within DruID. For this, we use a deep neural network (DNN). The DNN is trained to take as input a pair – drug and mutation profile – and predict the AUC for the drug on that profile. The mutation profiles are 324-dimensional binary vector representing the 324 genes in the FoundationOne CDx panel. A value of 1 in the *i*^th^ position indicates a mutation in the *i*^th^ gene. The drugs are represented by 2048-dimensional Morgan fingerprint bit vectors that encode the chemical structure. Thus, the input to the DNN is a (2048 + 324 =) 2372-dimensional binary vector and the output is a real number indicating the AUC.

Our DNN has a feedforward network architecture comprising 3 layers. The first two layers are of size 1000 and 100 with Tanh and ReLU activations. The final output layer has a sigmoid activation to obtain an output value that lies between 0 and 1 to represent the AUC. The network is trained to minimize the mean squared error loss, using the Adam optimizer [41]. Dropout with rate 0.5 is used to prevent overfitting and batch size of 1000 is used. Training is done for 50 epochs. Note that DruID only utilizes the predictions of the model. Any other predictive model, e.g., DrugCell [25], may also be used within DruID.

**Figure 1:**
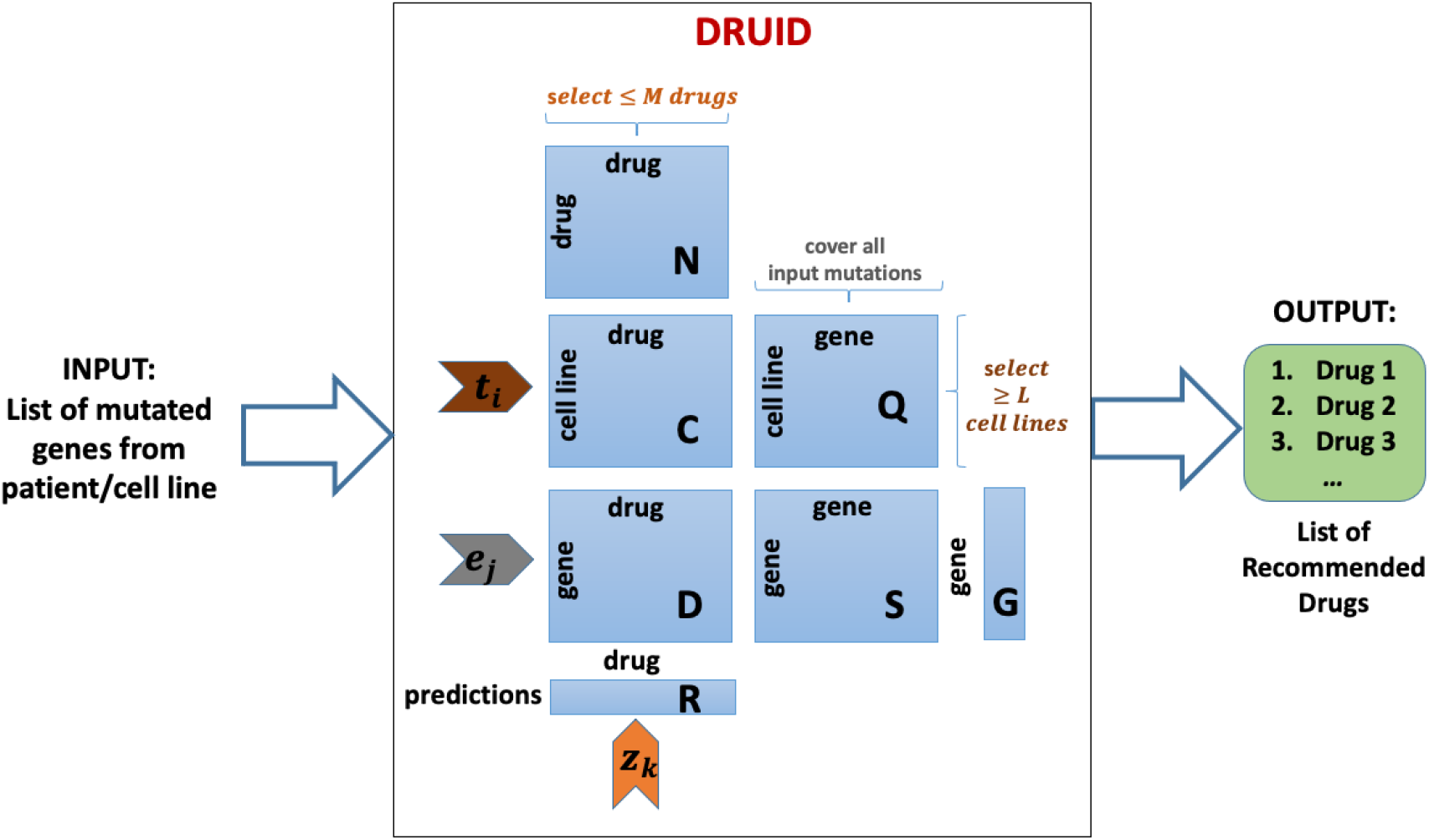
A schematic of DruID. Decision variables are *t_i_,e_j_,z_k_* that are used to select cell lines, genes and drugs respectively. *L* and *M* are user inputs that affect the number of cell lines and drugs selected. Other constraints are described in the text.

## 3 Empirical Evaluation

We evaluate DruID using cell line data on which the efficacy of many drugs have been tested and thus provides us with “ground truth” of drug efficacy which would not be available for real patients. Each cell line comes from a different tumour and serves as a proxy for a real patient. Note that while other information about these cell lines could potentially be used, such as gene expression, we restrict ourselves to mutations from the cell lines to simulate real life settings of diagnostic panels that provide mutation information only.

Our input data consists of 980 cell lines from GDSC (also present in CCLE) on which sensitivity to up to 400 drugs have been evaluated. We consider 324 genes that are used in the FoundationOne CDx panel. For the 980 cell lines, we obtained the list of mutations in these 324 genes from CCLE. Both variant types ‘damaging’ and ‘other non-conserving’ mutations were included. We retrieved 1,461 synergistic and 15 antagonist drug interactions across these 400 drugs from DDI databases. Synthetic lethality (SL) interactions for the 324 genes were retrieved from SynLethDB. GScores for the 324 genes and DScores for all pairs of 400 drugs and 324 genes were obtained from PanDrugs. We retrieved all 400 drugs’ chemical structure from PubChem [42] in SMILES format and then used the RDKit open-source cheminformatics tool [43] to convert the SMILE chemical structure into 2,048 Morgan fingerprint bit-vector for each drug.

### 3.1 Experiment Setting

We conduct two experiments to evaluate DruID. First, we compare its performance with that of PanDrugs. Second, we investigate the effect of each objective term within DruID through an ablation study. This is conducted by comparing the performance of the model as presented above with models where each objective term is removed, one at a time. These terms include CD: Cell line Drug Efficacy term, GS: GScore term, SL: Synthetic Lethal Interaction term, DDI: Drug-Drug Interaction term, NN: Neural Network prediction term. Both the experiments are done on 3 randomly selected test sets from the cell line data, by dividing the input 980 cell lines into train and test splits, consisting of 80% and 20% of the cell lines respectively. Cell lines with no mutations in the considered 324 genes were removed in each test set.

For each test cell lines GDSC has a different number of drugs, ranging from 123–395, whose sensitivity have been tested. Let *f_i_* denote the number of drugs whose AUC values are known, for the *i*^th^ cell line. For DruID, only the cell lines in the train set were considered in input matrices C and Q. All pairs of cell lines and the associated *f_i_* drugs were used to train the neural network. The predictions of this model was used to obtain vector R of predicted drug efficacies for each cell line in the test set. DruID was run, with *α* = 1, *β* = 5, for *M* = 5, 10, 15, 20 to obtain four different lists of drug recommendations. For each input cell line in the test set, we randomly select cell lines in the train set until the selected cell lines contain all input mutations. We set *L* as the number of cell lines selected. This approximates the minimum number of cell lines covering all input mutations and is sufficient since the optimization within DruID can select more cell lines if required as *L* is a lower bound.

To obtain the performance of PanDrugs, for each test cell line we input the list of mutations (from the 324 considered mutations) to PanDrugs and obtain their recommendations using their backend API [44], where we set the biomarker and direct target gene parameters to be true. Their recommendations are sorted in descending order of DScore. For each value of *M*, we consider the top 5, 10, 15 and 20 drug recommendations respectively.

Let *A_ik_* be true value of the efficacy of the *k*^th^ drug on the *i*^th^ cell line (from GDSC). Let 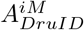 and 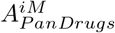 be the mean efficacy of the recommended *M* drugs by DruID and PanDrugs respectively. The mean efficacy is given by the average (true) efficacy of recommended drug on the *i*^th^ cell line: 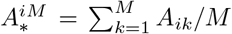, where * indicates DruID or PanDrugs. For each test set, we report the average of the mean efficacy for both the methods 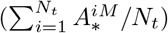, where *N_t_* is the number of cell lines in the test set.

### 3.2 Results

Table 2 shows the mean efficacy of DruID and PanDrugs over 3 randomly selected sets of cell lines, for *M* = 5, 10, 15, 20. Note that smaller AUC values indicates better drug efficacy, and so, lower values indicate better performance. Our results show that the integrative approach of DruID outperforms PanDrugs. For all four values of *M*, the drugs recommended by DruID had better efficacy (lower AUC) in nearly all cell lines.

**Table 2:**
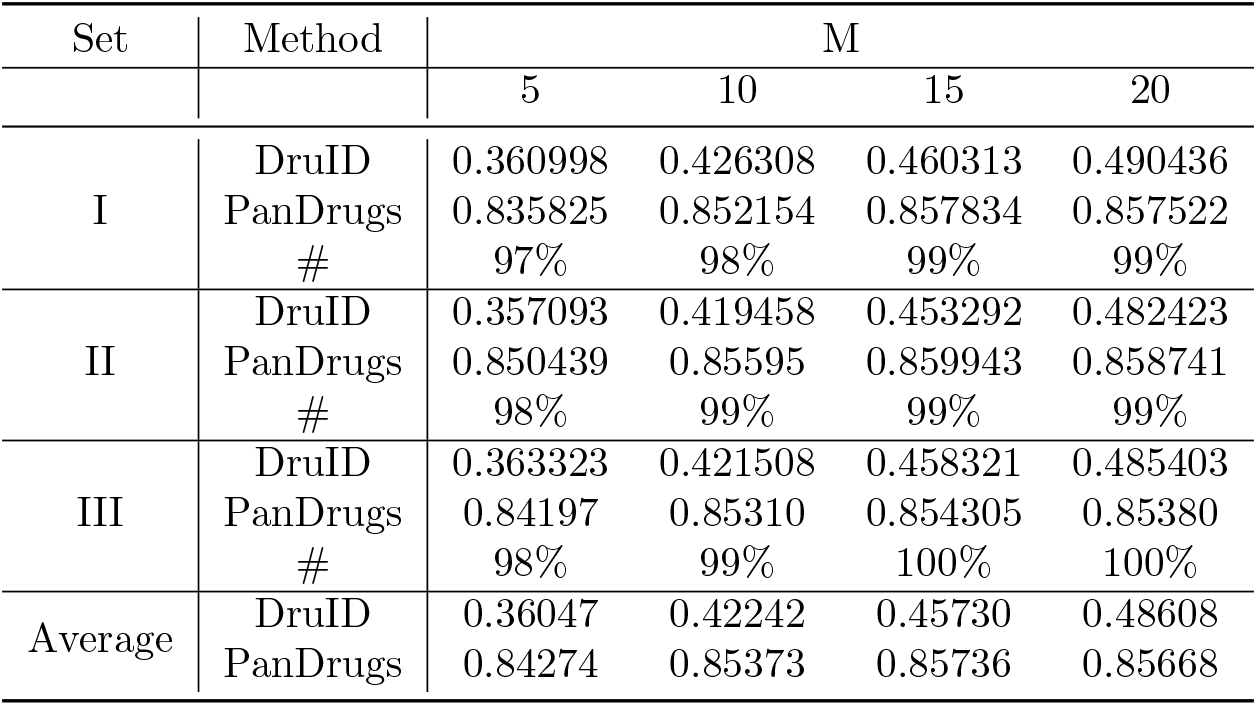
Mean efficacy (lower is better) obtained by DruID and PanDrugs in 3 randomly selected test sets. # indicates percentage of cell lines on which DruID obtains better AUC compared to PanDrugs.

Table 3 shows the results of our ablation studies. The lowest (best) mean efficacy values, for *M* = 20, are obtained by DruID. Removing the objective function terms corresponding to CD, GS, DDI and NN within DruID increases the AUC (lowers the performance). This shows that each of these terms contributes to the performance of DruID. No SL partners were found for these input genes in the data we considered, and so the SL term (and its removal) had no effect on the optimization. Further, the neural network alone also obtains AUC that is higher (lowers the performance) than that of DruID. This indicates that combining neural network predictions within the ILP framework of DruID helps in improving the overall performance.

**Table 3:** Comparison of DruID, Neural Network (NN) within DruID and component-wise removal of objective terms within DruID. CD: Cell line Drug Efficacy term, GS: GScore term, SL: Synthetic Lethal Interaction term, DDI: Drug-Drug Interaction term, NN: Neural Network prediction term. All results are for M=20.

| Set | DruID | [DruID - CD] | [DruID - GS] | [DruID - SL] | [DruID - DDI] | [DruID - NN] | NN |
| --- | --- | --- | --- | --- | --- | --- | --- |
| I | 0.490436 | 0.532594 | 0.533016 | 0.490436 | 0.52658 | 0.498091 | 0.496963 |
| II | 0.482423 | 0.540918 | 0.541149 | 0.482423 | 0.530251 | 0.482423 | 0.489172 |
| III | 0.485403 | 0.540241 | 0.539433 | 0.485403 | 0.530468 | 0.491612 | 0.495418 |
| Average | 0.486087 | 0.537917 | 0.537866 | 0.486087 | 0.529099 | 0.490708 | 0.493851 |

## 4 Discussion and Conclusion

In this paper we present DruID, an optimization framework that internally utilizes a deep neural network, and combines its predictions with a variety of evidence from public databases related to multi-gene markers, drug response screens, gene essentiality and clinical status of drugs, among others. Our empirical results show that DruID can effectively integrate multiple sources of information and find potentially useful drug combinations for a patient. Its performance is superior to that of PanDrugs, the best previous method designed for the same purpose. Both PanDrugs and DruID use GScores and DScores. The difference lies in the way they are used to prioritize drugs. Moreover, DruID integrates predictions from a deep neural network and information on drug-drug interactions and synthetic lethal interactions that can potentially improve its final recommendations both with respect to drug efficacy and choices provided to the clinician.

The genes we considered in our experiments, both for the predictive model as well as in cell lines were restricted to 324 genes from the Foundation One report. This was a deliberate choice to mimic the practical setting where clinicians have information of these genes only from the diagnostic panel report. Our framework is not limited to this setting, and can consider other genes as well. DruID is also not limited to the neural network we used in our experiments; any other predictive model may be used including those that offer more interpretability to the clinician, such as decision trees or logistic regression.

Our framework can be modified and/or extended to consider other kinds of data by adding appropriate terms and constraints to the optimization model. Integrating functional dependencies, in addition to the considered SL interactions, protein interaction networks, pathway contexts, transcriptional regulatory modules and clinical data may improve the modeling of latent correlations and help in exploring alternative treatments. Our model may also be made more specific with respect to selection of cell lines. For instance, the alterations given in the diagnostic panel may be reverse annotated [45] to match the exact genomic variant in the cell lines. This would make the matching more stringent and reduce the number of cell lines, and thus may be considered when more cell line data is available. The use of multiple data sources and models within DruID may allow new ways of interpreting the evidence for decision making. Future work can explore these research directions. It would also be interesting to evaluate the utility of DruID in a real clinical setting.

We assert that the aim of DruID is not to replace the clinician but only to provide decision support. The amount and heterogeneity of evidence available from various sources that can potentially aid the decision of which drugs to combine and prescribe is increasing rapidly. DruID aims to integrate such information from multiple databases and present it to the clinician for subsequent decision-making. Our hope is that the efficacy of integrative methods like DruID would continue to improve with increasing availability of high resolution genomic data in public databases, to the benefit of the prescribing clinician.

## Notes

### Competing Interest Statement

The authors have declared no competing interest.

